# Implementation and calibration of the Vaganov-Shashkin model in the virtualRings R package

**DOI:** 10.64898/2026.09.01.748354

**Authors:** Feng Wang, Jeff W. Atkins, Kevin J. Anchukaitis, Erika K. Wise, Xiuchen Jiang, Bao Yang, Dominique Arseneault, Étienne Boucher, Matthew P. Dannenberg

## Abstract

Process-based tree growth models provide a mechanistic framework for investigating how climate conditions regulate tree growth across daily to annual time scales. Yet, their broader application across species and environments is constrained by the limited accessibility in open-source environments and the difficulty of estimating physiological parameters that are rarely measured directly. Here, we present virtualRings, a new R package integrating the Vaganov-Shashkin model (VSM) and the RINGS3 models, and focus on the implementation and calibration of VSM. Using tree-ring width observations from seven Northern Hemisphere sites across various environmental conditions, we compared the traditional bootstrap-based calibration approach with the Covariance Matrix Adaptation Evolution Strategy (CMA-ES). CMA-ES improved agreement between simulated and observed radial tree growth and provided an efficient approach for model parameter estimation. We further evaluated practical CMA-ES settings to balance computational cost and performance and discussed its potential limitations. The virtualRings package provides an open and reproducible platform for tree growth simulation, facilitating the application of important process-based models across species and environments and the investigation of how temperature and moisture constraints regulate daily tree-ring formation across spatial and temporal scales.

## 1. Introduction

Tree rings are widely used to study climate variability, forest ecology, and forest carbon sequestration. While external and internal factors regulate wood formation, the intra-and inter-annual structures of tree rings can, at least partially, reflect changes in climate and environment during tree growth (Schweingruber, 2012). Thus, tree rings offer invaluable insights into both past climate variability (Anchukaitis, 2017; St. George, 2014) and ecosystem responses to past, present, and future climate variability and change (Babst et al., 2018; Dannenberg et al., 2019). Most tree-ring analyses rely on data measured at annual time steps (e.g., ring width, wood density, and isotopes), surpassing many other paleoclimate proxies (e.g., sediments and speleothems; Christiansen & Ljunsgqvist, 2017) and ecological observations (e.g., inventory-based stem diameters (Evans et al., 2017)) in resolution. However, annual tree-ring data are insufficient to study tree growth dynamics at daily to seasonal scales, which are essential for understanding wood formation dynamics (Cuny et al., 2014; Cuny & Rathgeber, 2016; Rossi et al., 2008) and nonlinear and non-stationary tree growth response to climate (Anchukaitis et al., 2020; Wilmking et al., 2020). For example, the “divergence problem” (the unstable or shifting tree ring–climate sensitivity) questions the suitability of tree rings (especially ring width) for climate reconstructions in some temperature-limited regions like Alaska (D’Arrigo et al., 2008; Esper & Frank, 2009). The causes of this phenomenon remain widely discussed, largely because the interaction between climate and wood formation is little studied, especially in remote regions.

Process-based tree growth models, such as the well-established Vaganov-Shashkin model (VSM; Vaganov et al., 2006), incorporate empirical observations from various stages of wood formation and numerical representations of eco-physiological processes (Eckes-Shephard et al., 2022). Compared with tree physiology (e.g., dendrometer or xylogenesis) studies that are often constrained by limited sites and relatively short observation periods (Cruz-García et al., 2019; Cuny et al., 2014; Deslauriers et al., 2007; Morino et al., 2021; Rossi et al., 2008), many process-based models driven by climate inputs allow for the investigation of sub-seasonal growth-climate relationships and/or cambial phenology across broader spatiotemporal domains (Buttò et al., 2020; Tumajer et al., 2021; Tychkov et al., 2019; Zelenov et al., 2024). After calibrating with conventional annual tree-ring observations, these models can be used to investigate tree phenology (Tumajer et al., 2021; Yang et al., 2017) and predict tree growth under future climate scenarios (Tumajer et al., 2025; Wise & Dannenberg, 2022). For example, using a simplified Vaganov–Shashkin model (Tolwinski-Ward et al., 2011), a recent study shows that future extensions of the growing season are unlikely to offset growth declines in European drought-prone forests (Tumajer et al., 2025). In addition, process-based models incorporating climate components can be used to reconstruct past climate via inverse modeling, although it can be computationally expensive and has been applied in limited cases (Boucher et al., 2014).

However, the implementation of process-based tree-ring models faces several challenges. First, different models were developed using programming languages with varying levels of complexity, thus requiring sufficient coding skills for effective and correct use (see summary in Eckes-Shephard et al., 2022). Models in compiled languages tend to be less flexible for updates, modifications, and enhancements compared with interpreted languages like R and Python, which have become increasingly popular in scientific data analysis (Atkins et al., 2022; Lortie, 2022). Although the computational time is slower in R and Python than in compiled languages, migrating commonly used process-based tree-ring models into open-source environments can lower the technical barrier, improve model transparency, and facilitate wider application and further model development.

Another challenge arises from model calibration, as process-based models can require a large set of poorly constrained parameters that determine the simulated eco-physiological processes of tree growth (Xue et al., 2026). Ideally, these model parameters (e.g., thermal and soil moisture thresholds for growth) should have ecological meaning in the real world (Tolwinski-Ward et al., 2013) and be measured empirically, but these observations are usually not directly available for most species at specific locations. Users must therefore either rely on parameters from the same or similar species at other sites (Friend et al., 2022) or, more frequently, parameter estimation by optimizing model outputs, e.g., comparing observed and simulated tree-ring width (He et al., 2019; Tychkov et al., 2019). Such parameterization approaches range from simple bootstrap-based searches within the plausible parameter space (Wise & Dannenberg, 2022) to more sophisticated Bayesian inference (Tolwinski-Ward et al., 2013), but this can be particularly challenging for complex models (Van der Meersch & Chuine, 2023).

Efforts have been made to facilitate the implementation of the VSM, the most frequently used climate-driven tree-ring model in dendrochronology (Eckes-Shephard et al., 2022). VSM uses daily temperature, precipitation, and latitude as input, and simulates tree-ring formation through nonlinear functions of temperature, precipitation, and day length (Vaganov et al., 2006). Earlier versions of the full VSM model were programmed in Fortran and Pascal (Anchukaitis et al., 2020), and their use was limited as a result. To mitigate obstacles in parameter estimation, an interactive, oscilloscope-like tool called VS-Oscilloscope was developed (Peresunko, 2018; Shishov et al., 2016). This user-friendly platform allowed for manual tuning of model parameters to obtain an optimal calibration, and thus is widely used (Gao et al., 2019; He et al., 2017; Tychkov et al., 2019; R. Xue et al., 2022). However, the manual nature makes VS-Oscilloscope difficult to apply across networks of many sites or species and to explore the full parameter space compared with automated approaches. A recent MATLAB version of VSM (Anchukaitis et al., 2020) improved the model’s accessibility (Tumajer et al., 2023; Wise & Dannenberg, 2022; H. Xue et al., 2023), but requires a MATLAB license and depends on user-defined parameters or calibration approaches (Wise & Dannenberg, 2022; H. Xue et al., 2023). Although the bootstrap method is more suitable for large-scale studies than the manual VS-Oscilloscope, it remains insufficient to thoroughly explore the high-dimensional parameter space for robust model calibration. Simplified versions of VSM (Tolwinski-Ward et al., 2011) reduce the number of model parameters, which makes parameter estimation possible using Bayesian optimization, yet applying such Bayesian optimization to the several dozen parameters of the full VSM remains computationally challenging.

In response to these challenges, here we present the virtualRings R package, which includes two important process-based tree growth models in this open-source interpretive language: the widely used VSM for simulating ring width (Anchukaitis et al., 2020), and a more recent RINGS3 model for simulating ring width and wood density (Friend et al., 2022). This package provides a new framework for input preparation, sensitivity analysis, visualization, and two statistical approaches for parameter estimation (or model calibration). This solves the two main issues that currently limit widespread use of process-based tree modeling: lack of availability in open-source interpreted languages and lack of automated model calibration approaches that are built into the model workflow. In this paper, we focus on the VSM growth model to illustrate the package’s overall workflow and to compare the calibration performance for the traditional bootstrap-based approach (BOOT) and a more flexible Covariance Matrix Adaptation Evolution Strategy (CMA-ES) optimization (Hansen, 2006; Hansen & Ostermeier, 2001), which has recently been applied to calibrate process-based species distribution (Van der Meersch et al., 2025; Van der Meersch & Chuine, 2023) and carbon cycle models (Lee et al., 2025).

## 2. Package workflow for VSM

virtualRings includes functions for data preparation, model implementation, and post-processing and visualization (Figure 1). The VSM implementation in virtualRings is largely migrated from its MATLAB version (Anchukaitis et al., 2020) and uses daily temperature, daily precipitation, and site latitude to simulate three daily growth rates from temperature, soil moisture, and day length in the Environmental block. The temperature and moisture growth rates *G_T_*(*t*) and *G*_W_(*t*) on day *t* are determined by a piecewise linear function, which uses four thresholds (minimum, optimal lower, optimal upper, and maximum; Supplementary Figure S1). The overall growth rate *G*(*t*) is then determined using Eq. 1:

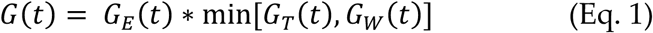

**Figure 1.**
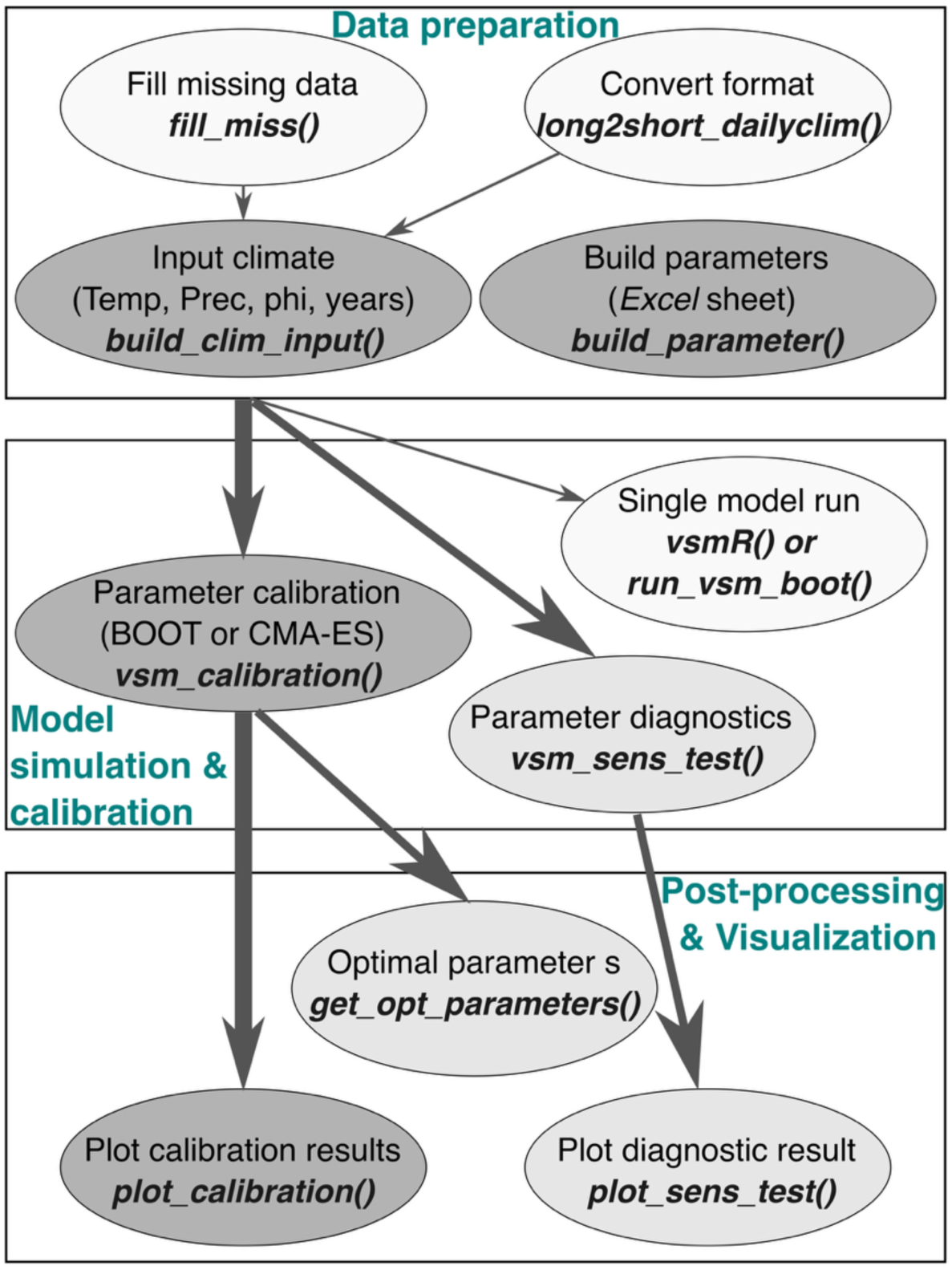
Workflow and key functions for running and calibrating VSM in virtualRings. The darkness of filled colors and the arrow width represent the potential importance and frequency of function usage (darker colors or thicker arrows: more important and frequently used).

where *G_E_*(*t*) denotes the rate due to day length. Subsequently, *G*(*t*) is used to indicate cellular growth rate in the Cambial block. Notably, like the MATLAB version, the current VSM in the package does not incorporate the Cell-size block, which simulates the cell features (i.e., dimensions and cell wall thickness) but requires substantial and site-specific parameters (Vaganov et al., 2006, 2011).

Here, we use the White Mountains (WHMT) tree-ring width chronology (Salzer et al., 2009) provided in the package (namely wmcrn) to demonstrate the main workflow and functionality for VSM. The WHMT chronology spans only 22 years (1956–1977), but it was one of the chronologies used to validate the MATLAB version of VSM (Anchukaitis et al., 2020), and is thus ideal for benchmarking our package. Daily climate data used for simulating the WHMT ring width are obtained from the ERA5-land dataset (the closest 0.1° grid to the site) (Muñoz-Sabater et al., 2021), and these data are also contained in the Package (i.e., wmclim). They can be loaded into the R environment using:

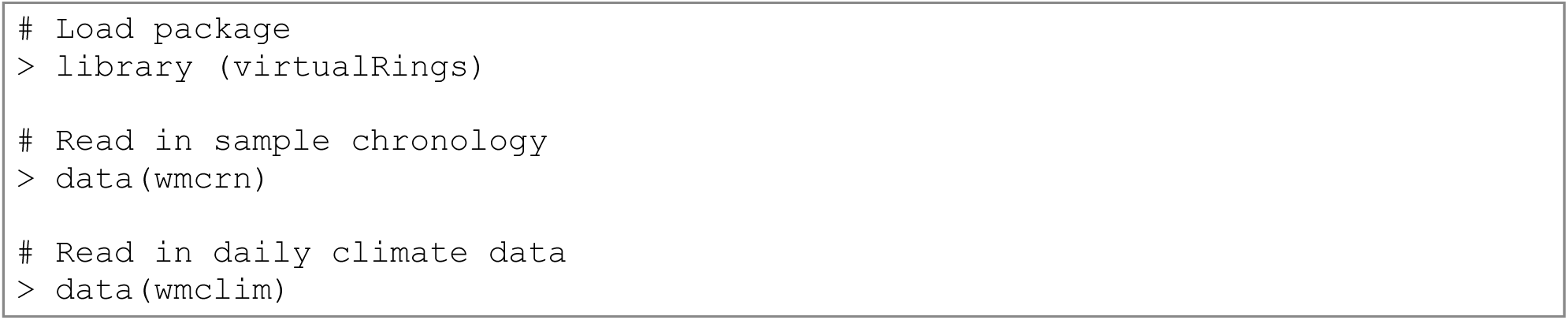

### 2.1 Data preparation

In the preparation stage, users generate a clim_input object with the build_clim_input() function, which can be directly used in subsequent model calibration and sensitivity testing functions. Daily temperature and precipitation data must be provided as a matrix with dimensions 366 × *M*, where *M* is the number of simulation years (eyear – syear + 1, where eyear is the end date and syear is the start date) and 366 represents the total number of days per year including leap years. The package also includes the function long2short_dailyclim() to convert three-column long format (“Year”, “day of year”, “climate data”, as it was in the original Fortran implementation), to a standard 366 × *M* data frame. Additionally, missing temperature values may be filled using clim_fillmiss(), which employs three gap-filling options (i.e., approximation, spline, or zero replacement). Temperature series may be filtered to mimic the smoothing effect of cambial temperatures (Edwards et al., 2021) using the same original Fortran filters as provided in VSM’s MATLAB version (Anchukaitis et al., 2020).

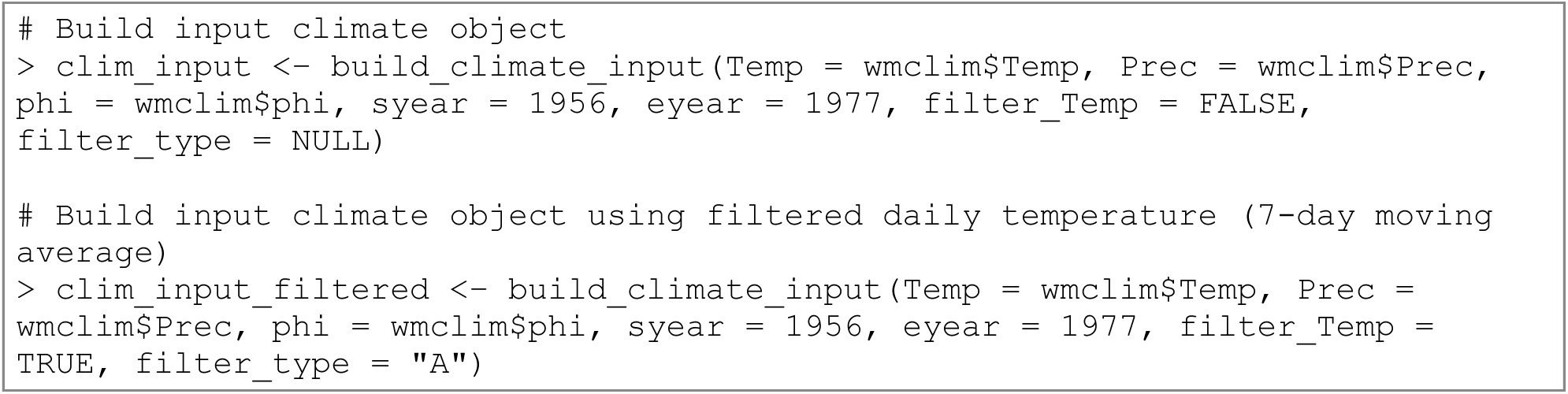

Default model parameters (Supplementary Table S1) and bounds can be defined in the VSM_parameter_sheet.csv file located in “inst/extdata”. While the values supplied with the package generally performed well in our experiments, it is possible to change the lower and upper parameter bounds, and if needed, the default value for each parameter in future applications. Other entries in the data sheet should remain unchanged to ensure that it can be used directly to construct the parameter input. When the lower and upper bounds are set to the same value for a given parameter, that parameter is treated as fixed, and therefore is not estimated during subsequent model calibration steps (Section 2.2). Parameter sets can be constructed in three ways: (1) boot for BOOT model calibration that randomly generates nboot sets of parameters (nboot = 1 for using default parameter values to run a single VSM), (2) cmaes for CMA-ES model calibration, and (3) stest for the one-at-a-time parameter diagnostics (sensitivity testing) by varying a single parameter while fixing all other parameters at their default values.

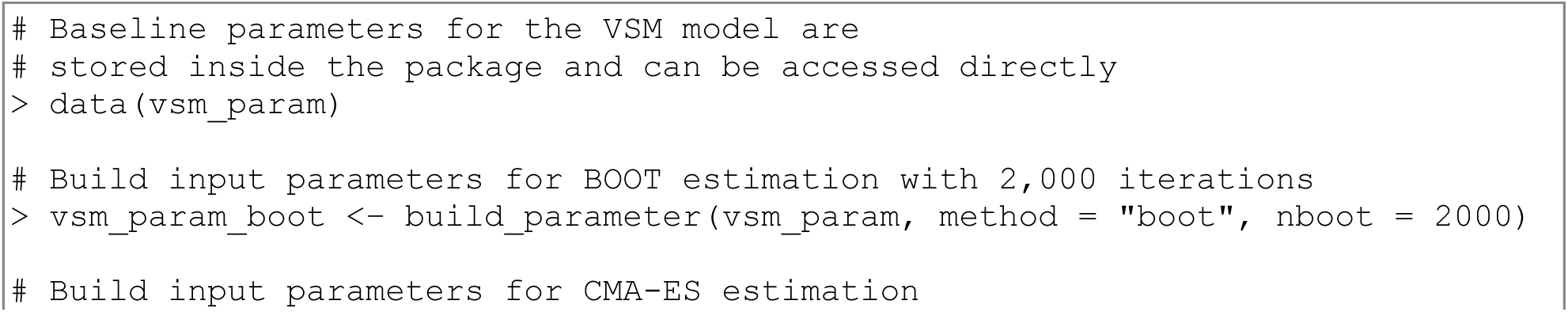
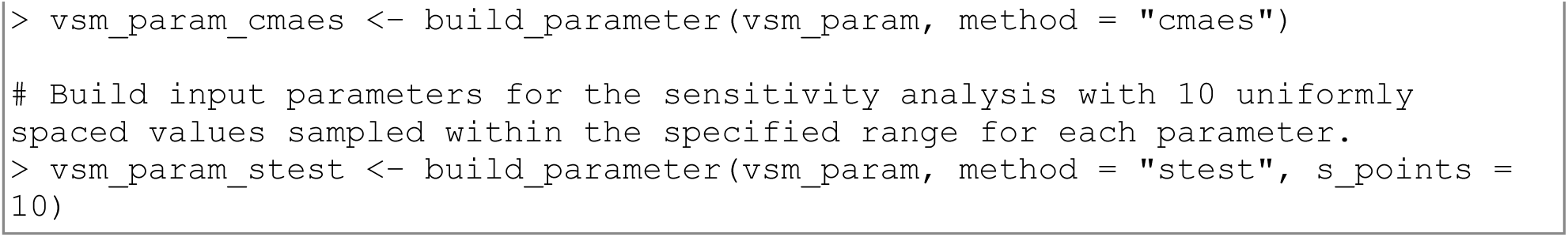

### 2.2 Model simulation, model calibration, and sensitivity testing

The package contains several key functions for model simulation and calibration. Although VSM can be run using a single or multiple sets of parameters (using vsmR() and run_vsm_boot(); see the package vignette), it is recommended to begin with the vsm_calibration() function to determine the optimal parameter estimates. As described previously, users may choose the BOOT or CMA-ES methods for model calibration, each of which uses the corresponding parameter list constructed with build_parameter(), along with the input climate data and annually resolved tree-ring observations. The observations typically consist of a detrended tree-ring chronology, in which age-related growth trends have been largely removed, because VSM does not incorporate the age effects of tree-ring formation. Since VSM returns two types of total ring width (TRW) indices (Anchukaitis et al., 2020)—one based on the normalizing number of xylem cells in each year (trw) and the other based on a normalized simulated cell size (trws)—the type of index needs to be specified using the argument tr_ind. The nbest argument specifies the number of best VSM simulations to retain, allowing for uncertainty evaluation of tuned model parameters obtained using the get_opt_parameter() function. In addition, the simulation performance can be evaluated using several different statistical metrics, including Root Mean Squared Error (stat = "rmse", default), Pearson’s correlation coefficient (stat = "r"), and the Kling-Gupta efficiency (stat = "kge") that jointly evaluates Pearson’s r and biases in mean and variation (Gupta et al., 2009).

Although methods are implemented within the same vsm_calibration() function, use of the CMA-ES method requires an additional cmaes_controls argument. The CMA-ES method in this package was primarily based on the algorithm modified by (Van der Meersch & Chuine, 2023), which itself is a modified version of the cmaes R package (Trautmann & Arnu, 2025). However, compared with the original CMA-ES algorithms that return a single best solution, our function can return an ensemble of the nbest VSM simulations.

Here, we briefly introduce the CMA-ES approach and the key cmaes_controls settings. This evolutionary algorithm starts with *λ* candidate solutions drawn from the initial multivariate parameter distribution and uses the best *μ* solutions (evaluated against observations) to update the distribution. Then, this procedure is iterated to find the best solution. In the cmaes_controls settings, *λ* and *μ* can be defined using lambda and mu, and sigma denotes the initial step size controlling the sampling distribution. The parameter maxit specifies the maximum number of iterations and serves as a stopping criterion. The scale_factor controls the linear rescaling of parameters to improve numerical stability during optimization. For example, all parameters are linearly scaled to the 0–10 range when scale_factor = 10, and we do not recommend modifying it. In addition, the CMA-ES convergence can be assessed using the stoptol argument. A value of stoptol = 0.1 indicates that the algorithm terminates when the range of each rescaled parameter falls below this threshold. The example scripts for using both calibration methods are shown below:

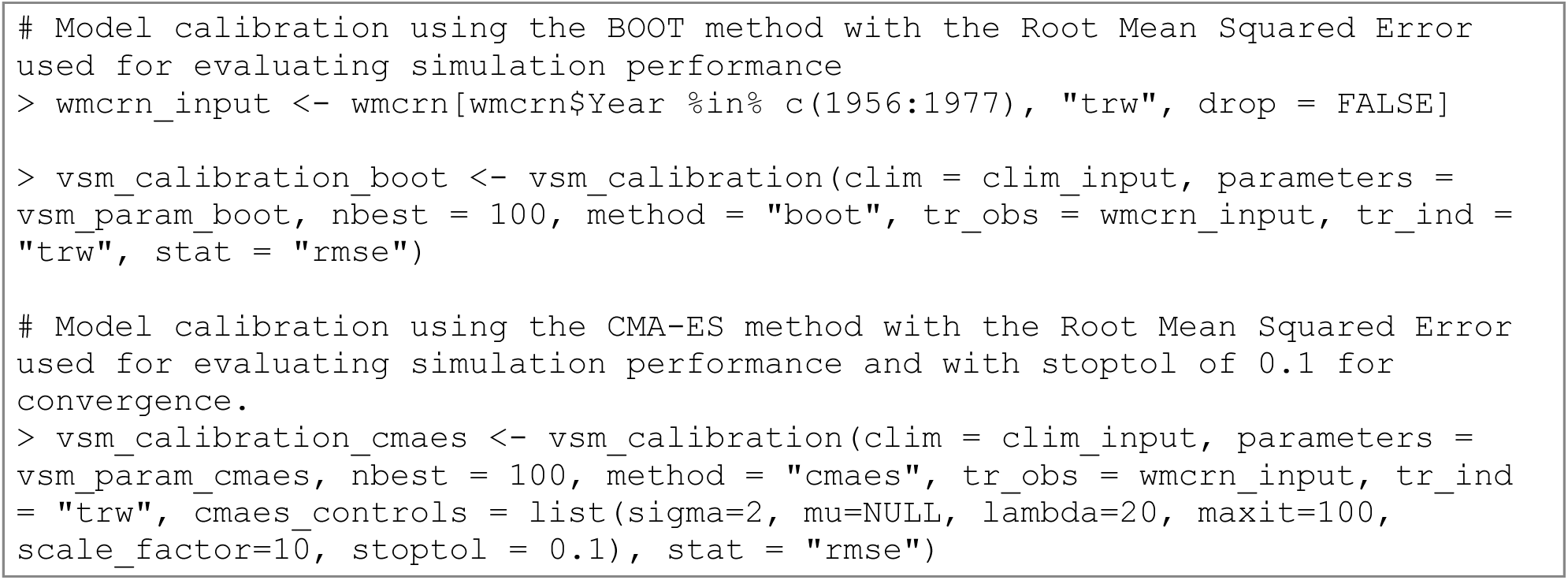

Sensitivity testing can be used to investigate the influence of individual parameters on model output. Although optional, this step is useful for reducing the dimensionality of the calibration problem and refining parameter ranges by identifying parameters to which the model is highly sensitive. Sensitivity analyses are typically “one-at-a-time” (Hamby, 1994) and are conducted by iteratively running the VSM while varying one parameter at a time and keeping all other parameters constant. It can be implemented using:

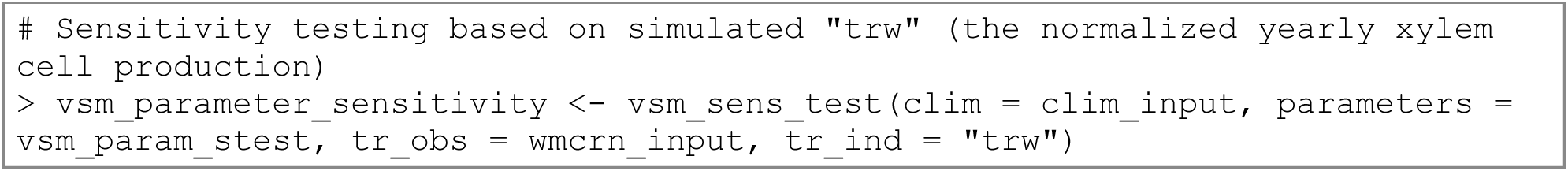

### 2.3 Visualization

virtualRings offers two functions to visualize the outputs of model calibration and sensitivity testing. The plot_calibration()function plots the comparison between the observed TRW and the nbest TRW simulations and, optionally, the range of calibrated model parameters (Supplementary Figure S2), while plot_sens_test() shows how a given validation metric varies across the specified range of each selected parameter (Supplementary Figure S3).

## 3. Comparison of calibration methods

### 3.1 Site and model settings

Since this represents the first application of CMA-ES for calibrating process-based tree-ring models like VSM, we compared its performance to the BOOT. We selected seven Northern Hemisphere tree-ring sites representing three different climate regions (Figure 2a) and tree species from three genera: *Pinus*, *Picea*, and *Cedrus* (Supplementary Table S2). Boreal sites include the Firth River in Alaska (FIRT, Anchukaitis et al., 2013; Andreu-Hayles et al., 2011; Edwards et al., 2025), L105 in Quebec (QUEB; Wang et al., 2020, 2022), and Tornesträsk in Scandinavia (TORN; Grudd, 2008). Tree growth at these sites is predominantly controlled by growing-season temperatures (Figure 2b), but TRW in Alaska is known for the “divergence” problem since the ∼1960s (D’Arrigo et al., 2008) and has a non-stationary association with climate over time (Andreu-Hayles et al., 2011). Attempts to create VSM simulations at FIRT have previously been unsuccessful, despite VSM’s ability to account for changes in growing limiting factors. For this site, we used an updated TRW chronology extending to 2022 (Edwards et al., 2025). The three arid or semi-arid sites include the Atlas Mountains in North Africa (ATAL; MORC019 in the International Tree-Ring Data Bank, accessed in July, 2025; Esper et al., 2007), Jingyuan in western China (JING; Yang et al., 2023), and WHMT in California (Salzer et al., 2009), all featuring limited precipitation and potentially warmer temperatures (Figure 2b). We also used a site in subtropical China, the West Tianmu Mountains (TIAN; Wang et al., 2019), where the radial growth of pine trees is affected primarily by temperatures during the summer and prior winter. Additional details are listed in Supplementary Table S1.

**Figure 2.**
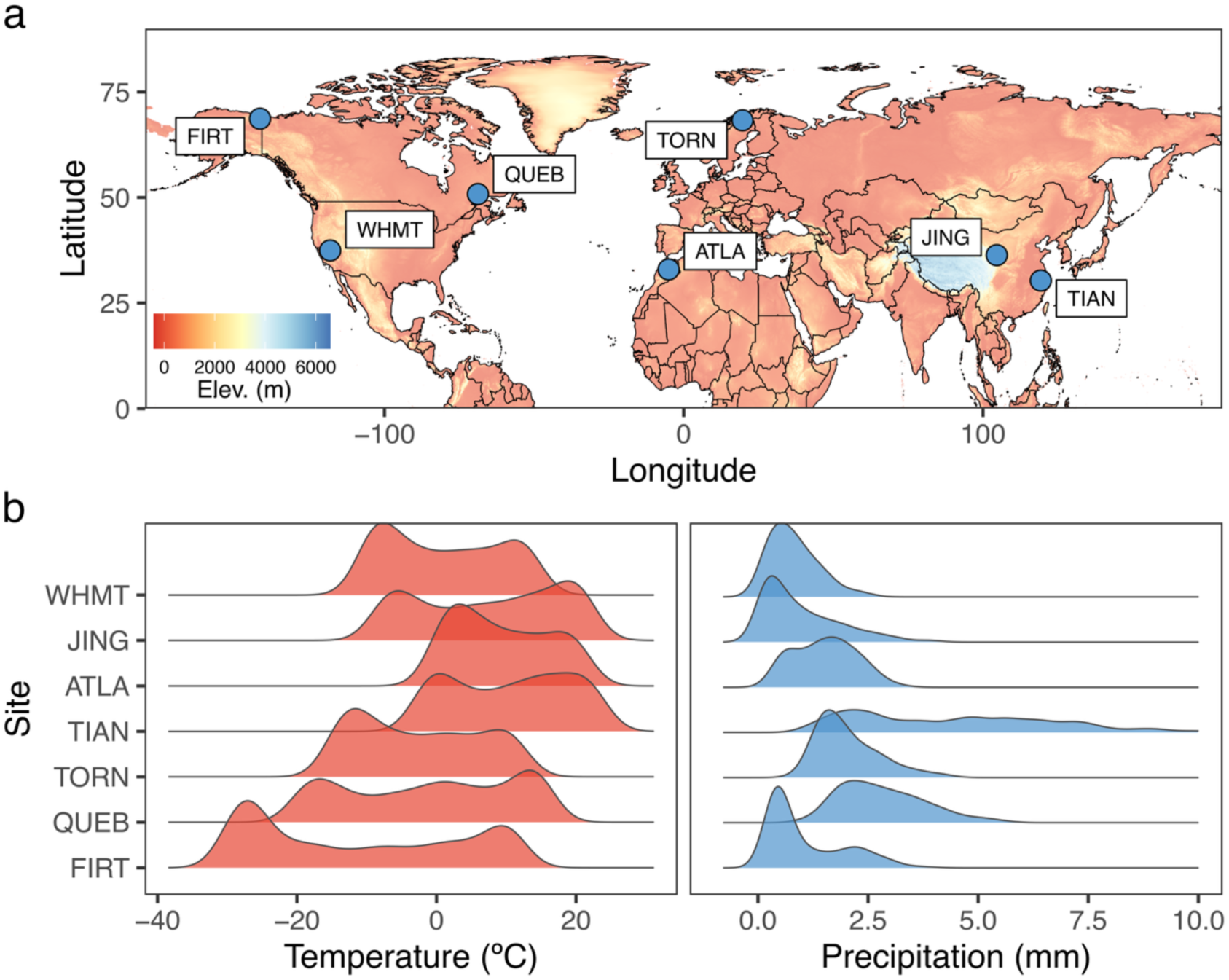
Location (a) and climate (b) of tree-ring sites used for VSM simulation. (b) shows the distributions of daily climate averaged over 1950–2023 at each site.

Daily temperature and precipitation data were obtained from the closest grid cells of the ERA5-land dataset. Although some differences remained when compared with *in situ* measurements, especially for precipitation, these biases had little effect on the interannual variability of simulated TRW (Wang et al., 2026). Considering the elevational differences between each sampling site and grid cell, temperature data were adjusted using a lapse rate of 0.6 °C per 100 m. This adjustment was only applied to sites showing elevational differences greater than 50 m, as the temperature biases would be minimal for sites with smaller elevational differences.

We followed the basic workflow of virtualRings for VSM calibration (Figure 1; Section 2). To compare the calibration performance of the BOOT and CMA-ES methods, the total number of iterations (*N*) was set to 1,000, 5,000, 10,000, 15,000, 20,000, 30,000, and 50,000 in seven calibration runs, and nbest was set to 100 to focus on the 100 best VSM simulation results (Section 2.2). The number of runs per CMA-ES iteration (*λ*) was set to 20, and the actual input maximum iteration (maxit) was set as 50, 250, 500, 750, 1,000, 1,500, and 2,500 for the CMA-ES runs (with no stoptol applied), given the relationship:

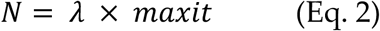

Additionally, we ran VSM calibration experiments with three stoptol values (0.1, 0.05, and 0.01) and *N* = 10,000 (maxit = 2,500, *λ* = 20) to investigate how stoptol settings affect the convergence of the CMA-ES algorithm. All calibrations were based on RMSE between simulated TRW (i.e., number of simulated xylem cells in each year divided by the mean cell numbers) and the corresponding TRW chronology. All computations were performed on a local machine equipped with an Apple M3 Max processor and 128 GB of RAM.

### 3.2 Simulated versus observed TRW

Figure 3 shows the consistency between simulated and observed TRW under different calibration settings for VSM. Overall, Pearson’s correlation coefficients increased with the total iteration number for both methods across the seven sites. However, Pearson’s *r* was approximately 0.2–0.5 greater when using the CMA-ES with *N* ≥ 5,000 (maxit ≥ 250) compared to the BOOT method. A total iteration number (*N*) between 5,000 and 10,000 yielded satisfactory and stable CMA-ES calibration performance, with a median *r* of 0.42–0.82 across the seven tested sites. This iteration (*N*) range generally aligns with calibrations using stoptol = 0.05 as the convergence criterion (Supplementary Figure S4), suggesting that the parameter search space becomes well constrained (i.e., an effective search tolerance of ∼0.05 relative to the 0–10 rescaled range) when maxit is set between 250 and 500, given *λ* = 20 (Eq. 2). Although the performance of the CMA-ES calibration may continue to increase slightly, a larger *N* value does not consistently result in a higher Pearson’s *r* (Figure 3).

**Figure 3.**
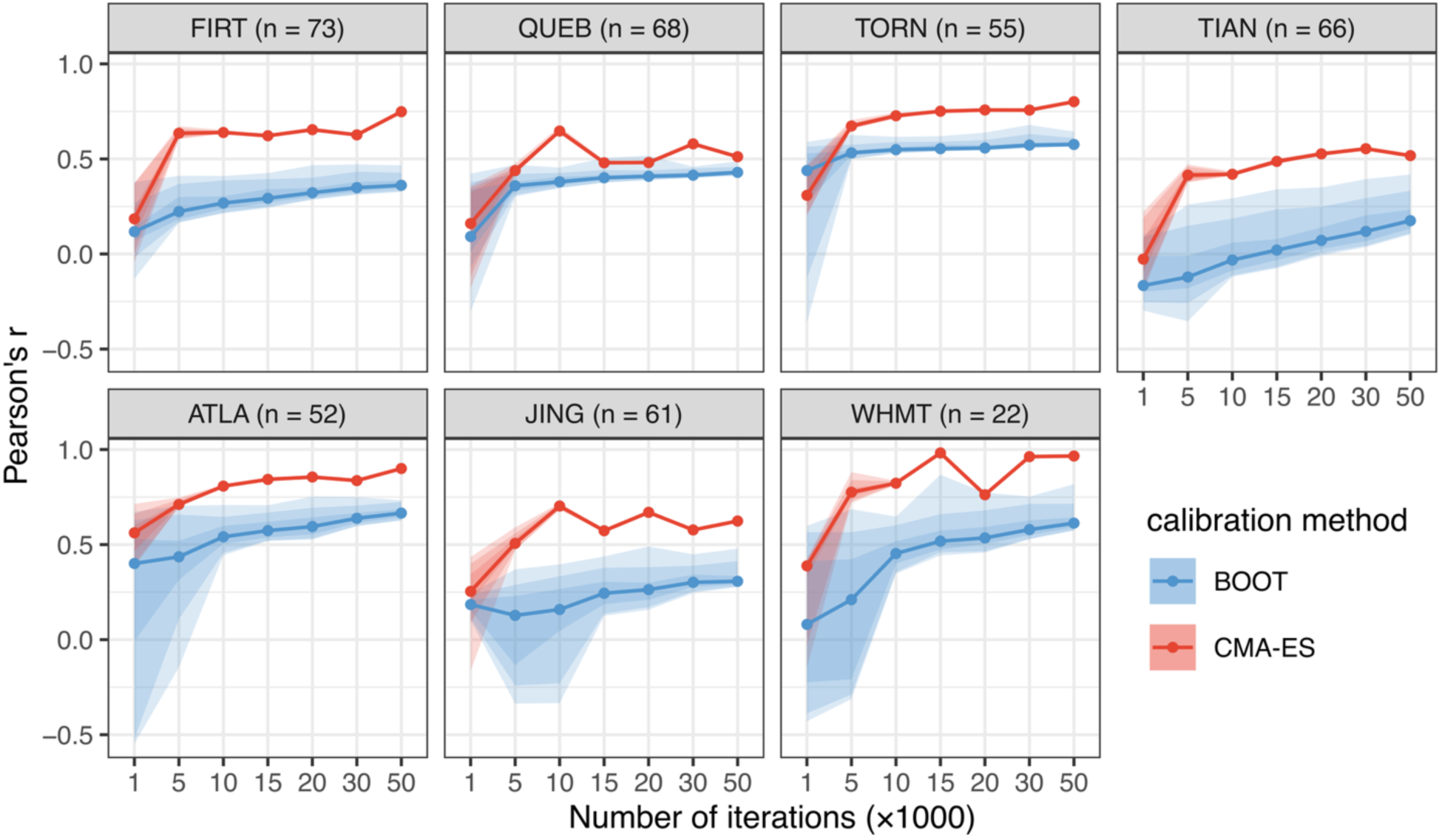
Pearson’s correlation coefficients between observed and the VSM simulated TRW using the BOOT and CMA-ES calibration methods and different numbers of iterations. Note that the number of iterations for the CMA-ES method equals a product of *λ* = 20 and *maxit* (Eq. 2). For example, a VSM run with an iteration of 10,000 means that *maxit* was set to 500, given *λ* = 20. Light to dark colored ribbons represent the ranges for the minimum to maximum, 5th–95th percentile, and 25th–75th percentile correlation coefficient.

In contrast, the BOOT method resulted in substantially weaker Pearson’s *r* values and wider spreads across the 100 best VSM solutions. Despite gradual improvements with increasing *N*, the BOOT experiments did not appear to reliably converge toward optimal VSM solutions. Even at *N* = 50,000, BOOT median correlations generally remain below those obtained using CMA-ES (Figure 3), and the 100 best simulated TRW series were less consistent, especially at sites FIRT, TIAN, and JING (Figure 4). These indicate that the BOOT method is less effective for solving the VSM calibration problem and would require more iterations to approach comparable performance, with no guarantee of stable convergence due to its dependence on random parameter sampling.

**Figure 4.**
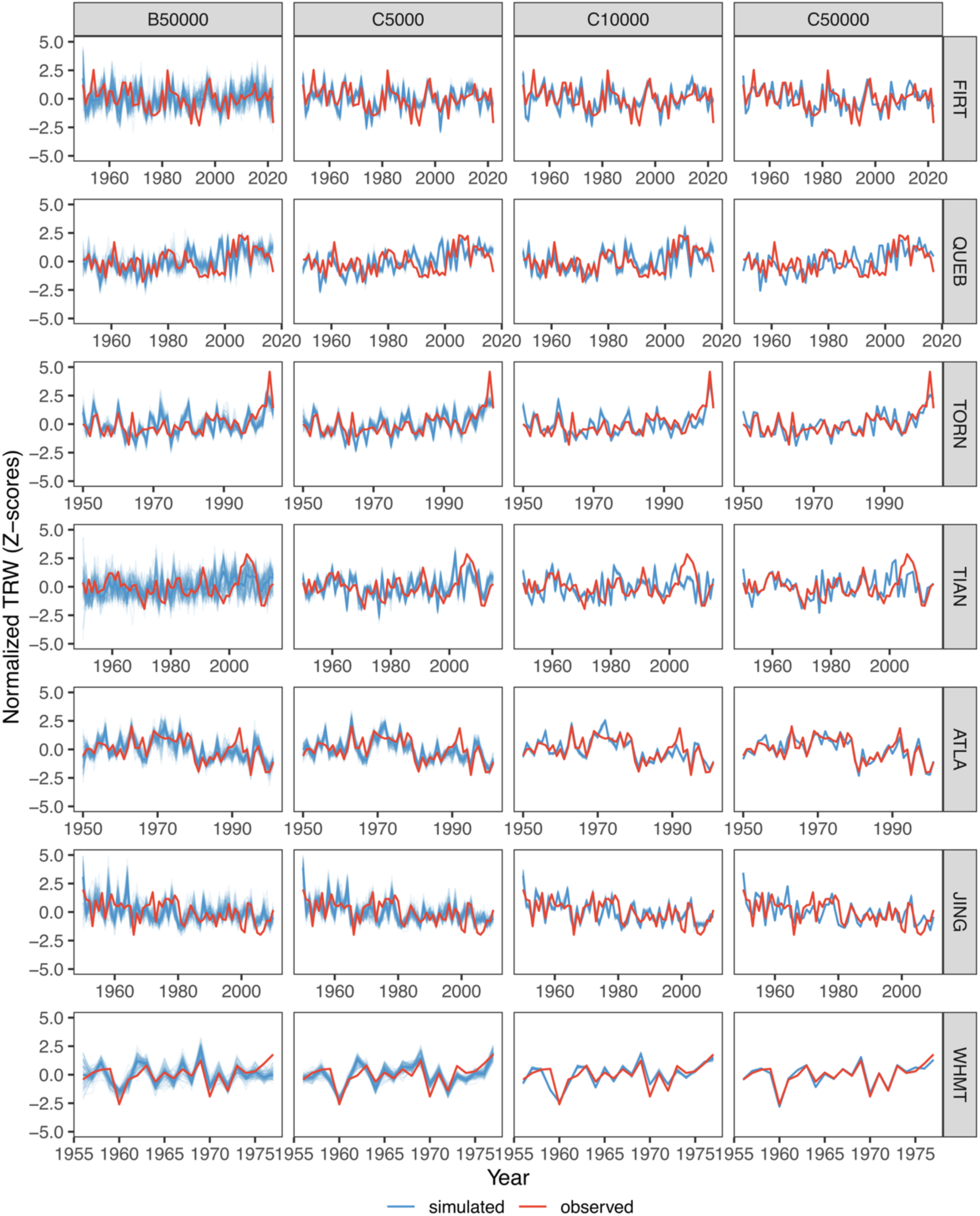
Comparison of the 100 best VSM-simulated (blue) and observed (red) TRW data for the 7 sites. B50000: the BOOT method with 50,000 iterations. C5000, C10000, and C50000: the CMA-ES method with 5,000 (*λ* = 20, *maxit* = 250), 10,000 (*λ* = 20, *maxit* = 500), and 50,000 (*λ* = 20, *maxit* = 2500) iterations, respectively.

### 3.3 Computation time

We further compared the computation time required for three CMA-ES calibrations with *N* = 5,000, 10,000, and 50,000 (hereafter referred to as C5000, C10000, and C50000), as well as the best-performing BOOT calibration (*N* = 50,000; B50000) (Table 1). Due to the different lengths of the seven ring-width chronologies (22 to 76 years), computation time was normalized to a uniform unit of 50 years to facilitate comparison. Specifically, the unit computation time was calculated by dividing the actual computation time by the corresponding chronology length and then multiplying by 50. The CMA-ES calibrations are computationally more expensive than the BOOT method. For example, for the same number of iterations (*N* = 50,000), the computation time was ∼10–20 times longer for the CMA-ES method. However, the C5000 calibration required a comparable computation time compared with the B50000 calibrations, while achieving substantially improved calibration performance at most sites (Table 1; Figure 4), with median *r* between simulated and observed ring-width chronologies ranging 0.41–0.78 for C5000 and 0.18–0.67 for B50000. These results point to the efficiency of the CMA-ES method in solving the parameter optimization problem for VSM.

**Table 1.** Unit computation time (hours per 50 years) and median Pearson’s *r* for 100 best simulations for selected VSM calibrated using the BOOT and CMA-ES methods. The computation time is based on a local machine equipped with an Apple M3 Max processor and 128 GB of RAM.

| Site ID | Unit computation time (hr / 50 yr) | | | | Median Pearson’s $r$ | | | |
| --- | --- | --- | --- | --- | --- | --- | --- | --- |
|  | B50000 | C5000 | C10000 | C50000 | B50000 | C5000 | C10000 | C50000 |
| FIRT | 0.68 | 0.84 | 1.76 | 5.90 | 0.36 | 0.64 | 0.64 | 0.75 |
| QUEB | 0.60 | 0.76 | 1.60 | 8.85 | 0.43 | 0.44 | 0.65 | 0.51 |
| TORN | 0.68 | 0.62 | 1.09 | 4.52 | 0.58 | 0.67 | 0.73 | 0.80 |
| ATLA | 0.59 | 0.68 | 1.99 | 11.96 | 0.67 | 0.71 | 0.81 | 0.90 |
| JING | 0.63 | 0.55 | 1.09 | 3.82 | 0.31 | 0.51 | 0.70 | 0.62 |
| WHIT | 0.65 | 0.99 | 2.51 | 13.74 | 0.61 | 0.78 | 0.82 | 0.97 |
| TIAN | 0.55 | 0.84 | 1.95 | 12.38 | 0.18 | 0.41 | 0.42 | 0.52 |

B50000: the BOOT method with 50,000 iterations. C5000, C10000, and C50000: the CMA-ES method with 5,000 ( $\lambda = 20$ , $maxit = 250$ ), 10,000 ( $\lambda = 20$ , $maxit = 500$ ), and 50,000 ( $\lambda = 20$ , $maxit = 2500$ ) iterations, respectively. Median Pearson’s $r$ is calculated from correlations between observed and the 100 best simulated ring-width series.

## 4. Discussions and Conclusion

In this paper, we introduce virtualRings, a new R package to address a long-standing need for an open-source, transparent framework for process-based tree-growth modeling. By integrating VSM and RINGS3 into a uniform R environment, the package provides a standard workflow for data preparation, model calibration, sensitivity testing, and visualization. Although we focus on the widely used VSM (Anchukaitis et al., 2020; M. N. Evans et al., 2006; Vaganov et al., 2006), a similar workflow (in vignettes) can be applied to the more recent RINGS3 model, which has the potential to model wood density driven by daily climate (Friend et al., 2022).

Specifically, we demonstrate the calibration of VSM using ring-width data from seven sites across the Northern Hemisphere, covering cold, arid, and humid ecosystems. The comparison between the two calibration methods implemented in virtualRings highlights the advantage of CMA-ES for parameter optimization in process-based models, as suggested by substantially improved correlations between simulated and observed TRW data relative to the conventional BOOT approach (Figure 3; Table 1). This improvement largely arises from the fact that CMA-ES updates the parameter search distribution based on the performance of a smaller set (determined by *λ*) of candidate solutions (Hansen & Ostermeier, 2001; Van der Meersch & Chuine, 2023). Consequently, parameter combinations that result in better objective-function values (i.e., evaluation scores against observations) are more likely to be retained and update subsequent searches, thus leading to more efficient convergence toward optimal solutions. In fact, CMA-ES performed particularly well at FIRT, where previous attempts to search for suitable VSM parameters using the BOOT method were largely unsuccessful (Figure 4). Most of the CMA-ES estimated parameters also fell within plausible ranges if using the default parameter values as a reference (Supplementary Table S3). In contrast, the BOOT method may provide limited coverage of possible combinations in the high-dimensional parameter space without adaptive updating of the search distribution.

According to our experiments using seven tree-ring chronologies, a setting of *λ* = 20 and maxit = 250–500 generations (equivalent to 5000–10,000 iterations) yields more robust calibration results than the BOOT method with 50,000 iterations, while requiring a comparable computational time. We therefore recommend a similar *λ* and maxit configuration for future CMA-ES-based VSM calibrations to achieve a good balance between optimization robustness and computational efficiency. Although a different *λ* value could be considered, we adopted *λ* = 20 following (Van der Meersch & Chuine, 2023) throughout this study. Theoretically, greater *λ* values tend to enhance the global search performance of CMA-ES, but they will require longer computational time, while smaller values can increase the risk of local convergence (Van der Meersch & Chuine, 2023). Higher generation numbers (e.g., maxit > 1,000) are not recommended as the CMA-ES algorithm may converge strongly, producing highly similar simulated TRW series (Figure 4).

Although powerful, CMA-ES may have limitations when calibrating process-based tree growth models. First, the risk of overfitting may increase when only a limited number of observations are available for calibration. This issue is especially relevant for tree-ring chronologies that have limited temporal overlap with station or gridded climate datasets that began in the early to mid-twentieth century (Menne et al., 2012; Wang et al., 2026). For example, the CMA-ES calibration at the WHMT site only spans 1956–1977, yet, Pearson’s correlations against the tree-ring chronology even exceeded 0.9 in some cases (Figure 3). Although these high correlations do not by themselves demonstrate overfitting, they suggest caution and the need to carefully evaluate the final parameters. Notably, all calibrations with extraordinarily high skill were characterized by substantially greater rooting depth and higher soil melting coefficient than calibrations showing lower but still reasonable skill (Supplementary Table S3). These rooting depths are probably unrealistic for bristlecone pine, which has a shallow root system (Fritts, 1969). These “problematic” parameters may therefore compensate for uncertainties in model simulations rather than accurately represent site conditions, reflecting coincidental agreement between simulations and observations. CMA-ES results should be interpreted with caution when the model is calibrated with relatively short tree-ring observations.

Second, CMA-ES may converge with different parameter combinations across different calibration runs. In our experiments, this is reflected in variations in calibration skill as the total number of iterations or maxit increases, in contrast to the progressive increases in skill when using the BOOT method (Figure 3). The variability among CMA-ES calibrations likely suggests both the potential equifinality of VSM and the stochastic nature of CMA-ES, which can lead to different runs with similar skill converging on substantially different regions of the parameter space. As a result, calibrations with comparable overall skill may yield different phenology-related VSM outputs (Supplementary Table S3, S4, Figure S5), which are widely used to investigate tree phenological responses to climate variability (Buttò et al., 2020; He et al., 2017; Tumajer et al., 2021; Tychkov et al., 2019; Yang et al., 2017; Zelenov et al., 2024). For instance, the minimum temperature threshold, Tf1, is clustered around 10 °C and 2.5 °C for the C5000 and C10000 calibrations at WHMT, respectively (Supplementary Figure S5; Table S3). The latter is closer to the widely observed critical temperature for tree stem growth across the Northern Hemisphere (Rossi et al., 2008). This problem of non-unique parameterization could be minimized by calibrating fewer parameters, for example, by conducting sensitivity tests and excluding parameters from calibration that exert minimal influence on results or by constraining the parameter space based on prior knowledge of plausible values for a given site or species. Although the BOOT method may be less sensitive to this issue due to its random-sampling nature, the spread of the selected phenological outputs under B50000 is substantially wider (Supplementary Figure S5), introducing greater uncertainties when interpreting tree phenology, such as the start and end of wood formation. Increasing maxit may further narrow the output uncertainties for BOOT calibration, but it will come at the expense of increased computational cost, and the convergence efficiency of BOOT has not been systematically evaluated in this study.

These results highlight the necessity of considering a set of optimal solutions when interpreting VSM-derived phenological patterns, regardless of the calibration method applied. Here, we propose several strategies to mitigate potential local convergence risks for CMA-ES:

1. A larger *λ* can be used to reduce the possibility of rapid local convergence; however, the computation time may increase.
2. A larger stoptol value (e.g., 0.1) can be used to permit a less conservative convergence criterion, thereby allowing a broader range of acceptable solutions to be retained.
3. Because relatively wide parameter bounds were used in our experiments, refining these ranges based on prior knowledge of tree phenology/physiology and results from sensitivity analyses may help reduce the risks of local convergence and improve calibration stability. Such refinement may also decrease the computational cost by limiting the exploration of implausible and meaningless parameter space.

In summary, this new virtualRings R package lowers the technical barrier to applying and, particularly, calibrating the widely used VSM process-based tree growth model. Importantly, calibration of physiologically meaningful parameters provides a means not only to improve tree-growth simulations but also to examine nonlinear climatic controls and diagnose thresholds governing cambial growth. For example, constraining parameters such as the minimum temperature threshold for growth initiation (Tf1) can help identify physiological limits on tree growth and examine how these controls vary across species and environmental gradients. The VSM implementation in this package is largely migrated from the MATLAB version (Anchukaitis et al., 2020) and therefore only contains the Environmental and Cambial block processes of the full VSM (Vaganov et al., 2011), while the heavily parameterized and species-and location-specific Cell-size block is not currently implemented. Nevertheless, the flexibility of the R environment allows for further model extensions, including the incorporation of additional growth modules and further model development, such as temperature-photoperiod interactions (Campelo & Camarero, 2024). In a companion paper (Wang et al., in prep.), we will demonstrate how VSM can be coupled with RINGS3 (Friend et al., 2022) to simulate climate constraints on wood density and evaluate RINGS3’s performance across multiple temperature-and drought-sensitive sites, extending beyond the original RINGS3’s application at a single high-latitude, temperature-limited site.

## Acknowledgements

This project was funded by the U.S. National Science Foundation P4CLIMATE program through grants AGS-2401699 (EKW) and AGS-2401700 (FW and MPD). FW also received support from a Haury Fellowship at the Laboratory of Tree-Ring Research, University of Arizona. KJA was supported by NSF grant 2124889.

## Author contribution

Conceptualization: FW, MPD, KJA, EKW. Methodology: FW, JWA, KJA, MPD. Software: FW, JWA, KJA. Formal Analysis: FW, XJ. Data Curation: FW, KJA, BY, DA, EB. Writing – Original Draft: FW. Writing – Review & Editing: FW, MPD, EKW, KJA, JWA, BY, DA, EB. Visualization: FW. Supervision: MPD, EKW, KJA. Funding acquisition: EKW, MPD.

## Data and code availability

Observed tree-ring-width data are available from the original references listed in Supplementary Table S2. The virtualRings R package is archived on Zenodo (https://doi.org/10.5281/zenodo.22116686) and is also available on GitHub (https://github.com/FengWang01/virtualRings).

