## supplemental for "Implementation and calibration of the Vaganov-Shashkin model in the virtualRings R package"

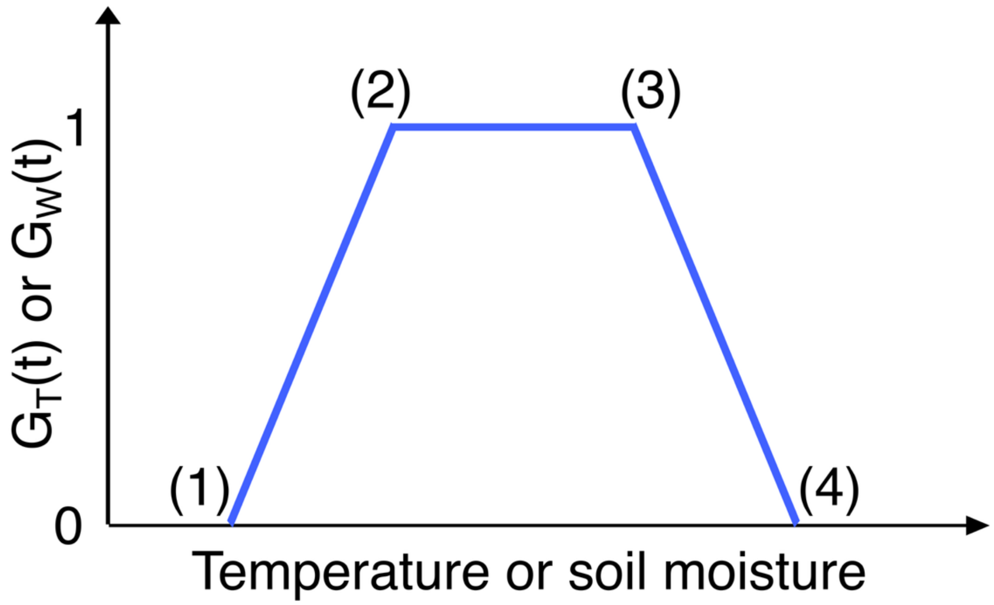


**Figure S1**. A piecewise linear function used to calculate daily growth rates dependent on temperature $G_T(t)$ and soil moisture $G_W(t)$. The four points (1)–(4) from left to right are the minimum, lower optimal, upper optimal, and maximum environmental thresholds.

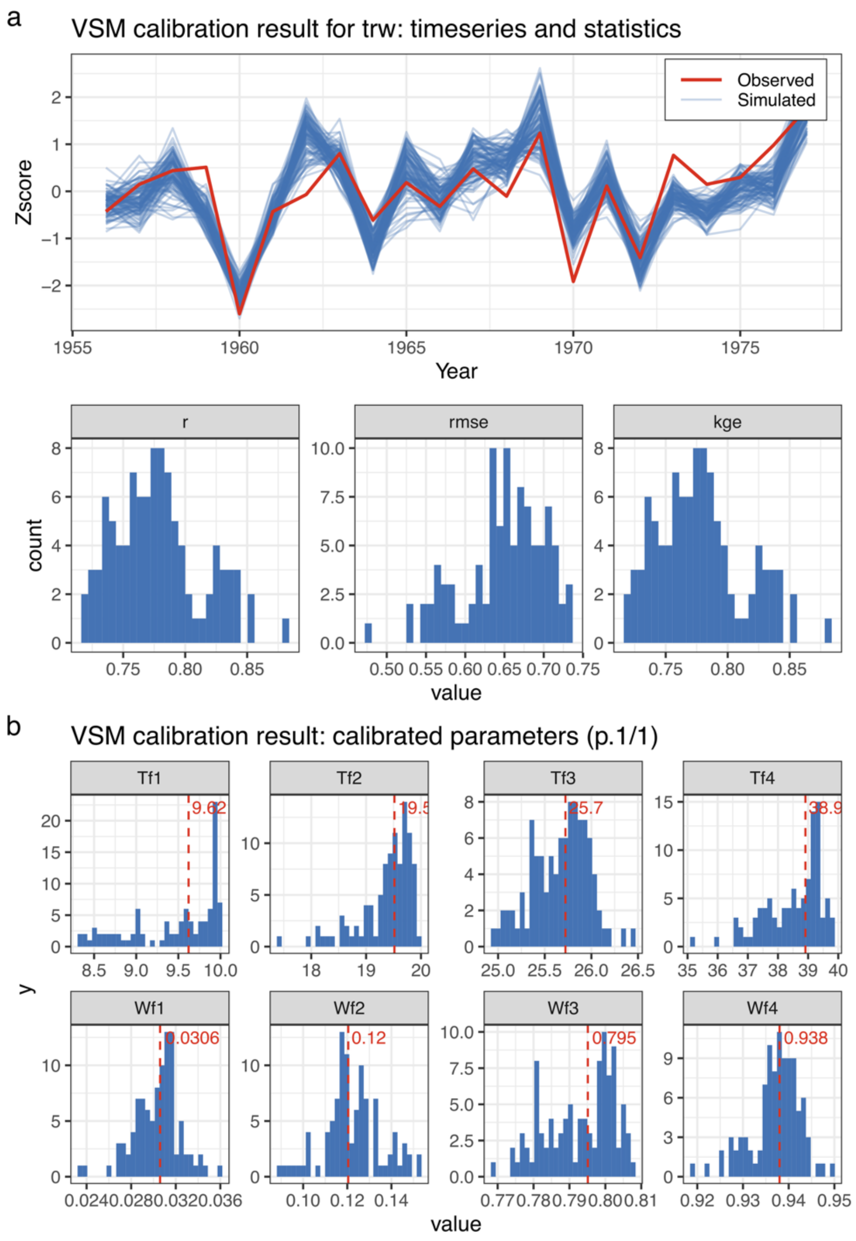


**Figure S2**. Visualization of calibration results using the `plot_calibration()` function. (a) Comparison between observed and 100 best-performed VSM simulations. The bottom panel in (a) shows the distribution of three statistical metrics between simulated and observed TRW data. r: Pearson's correlation coefficient; rmse: Root-Mean-Square-Error; kge: the Kling-Gupta efficiency. Because the statistics were based on Z-scored time series, kge is equivalent to Pearson's r. (b) Distribution of calibrated parameters for the `nbest` VSM simulations, with the red dashed line indicating the median (optional). The calibration results are based on the CMA-ES method with 5,000 iterations at WHMT.

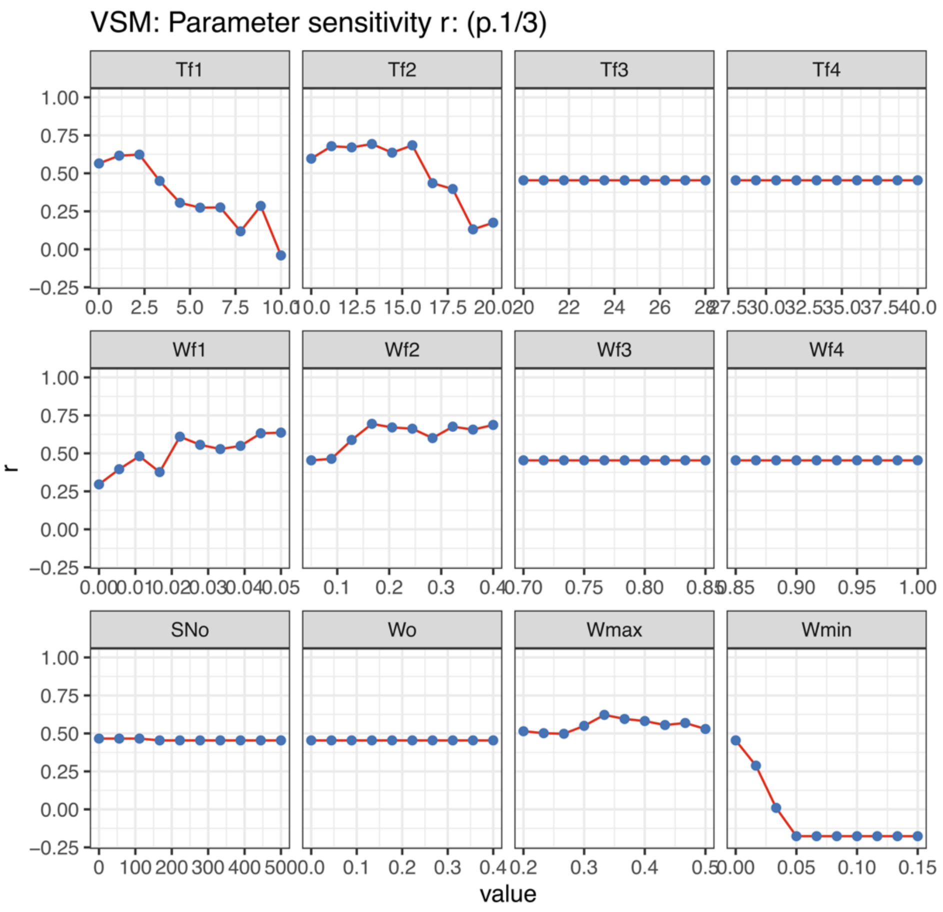


**Figure S3.** Visualization of sensitivity testing using the `plot_sens_test()` function. The results are based on the WHMT ring-width chronology. Each panel shows Pearson's *r* between the observed TRW chronology and the VSM simulated TRW series with one parameter varied (the x-axis) while all other parameters are fixed. Parameters (Supplementary Table S1) are displayed across three pages, with Page 1 (1/3) shown here.

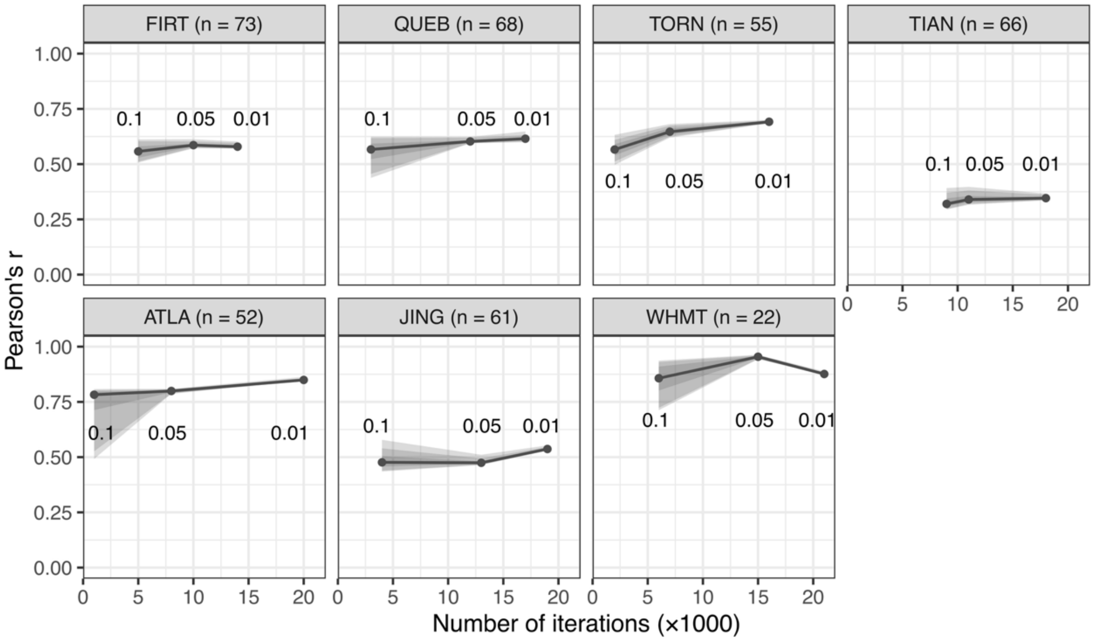


**Figure S4.** Same as in Figure 3, but for CMA-ES calibrations with different `stoptol` settings (0.1, 0.05, and 0.01) and a total iteration number ($N$) of 50,000. The X-axis shows the actual $N$ when the algorithm stopped.

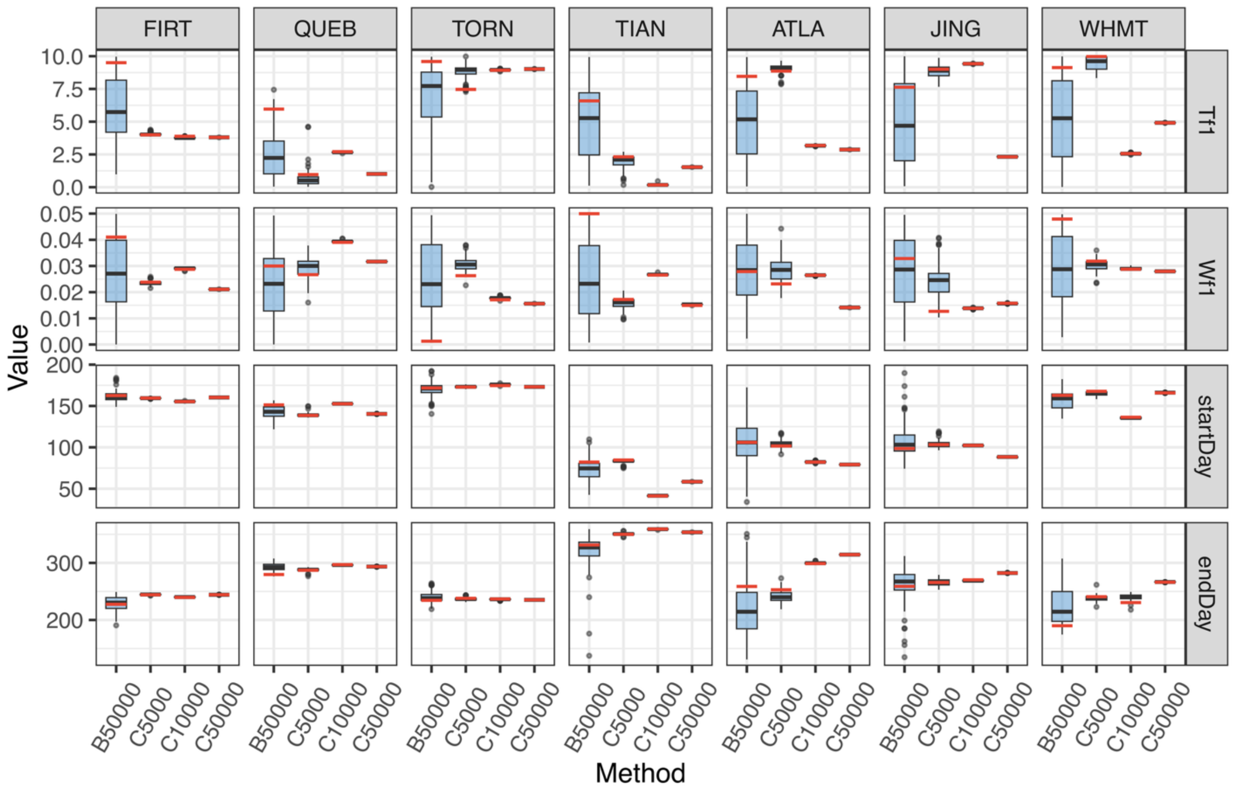


**Figure S5.** Distribution range for the calibrated minimum optimal temperature (Tf1) and moisture (Wf1) as well as the output start and end days of cambial activity from 100 best VSM calibration results following the BOOT and CMA-ES methods. B50000: the bootstrapping method with 50,000 iterations. C5000, C10000, and C50000: the CMAES method with 5,000 ($\lambda$= 20, $maxit$ = 250), 10,000 ($\lambda$ = 20, $maxit$ = 500), and 50,000 ($\lambda$ = 20, $maxit$ = 2500) iterations, respectively. Box boundaries show the 1st and 3rd Interquartile Ranges (Q1 and Q3), and tails show 1.5 times Q1 and Q3. The thick black and red bars represent the median values and values for the best calibration, respectively.

Table S1. VSM parameters that need calibration.

| Symbol | Description (unit) | Default value |
|---|---|---|
| Tf1 | Minimum temperature for growth (ºC). | 3 |
| Tf2 | Lower optimal temperature (ºC). | 16 |
| Tf3 | Upper optimal temperature (ºC). | 22 |
| Tf4 | Maximum temperature for growth (ºC). | 30 |
| Wf1 | Minimum soil moisture for growth (v/v). | 0.01 |
| Wf2 | Lower optimal soil moisture (v/v). | 0.05 |
| Wf3 | Upper optimal soil moisture (v/v). | 0.8 |
| Wf4 | Maximum temperature for growth (v/v). | 0.9 |
| SNo | Initial snowpack (mm). | 300 |
| Wo | Initial soil moisture (v/v). | 0.2183 |
| Wmax | Maximum soil moisture (field capacity, v/v). | 0.7 |
| Wmin | Minimum soil moisture (wilting point, v/v). | 0.01 |
| rootd | Root/soil melt depth (mm). | 500 |
| rated | The rate of water drainage from soil. | 0.0051 |
| Pmax | Maximum rate of infiltration water into soil (mm/day). | 30 |
| k1 | The interception precipitation by the tree crown. | 0.9 |
| k2 | The 1st coefficient for calculation the transpiration. | 0.25 |
| k3 | The 2nd coefficient for calculation the transpiration. | 0.18 |
| Tm | Sum of temperature to start soil melting (ºC). | 20 |
| Tg | Sum of temperature to start tree growth (ºC). | 30 |
| a1 | The 1st coefficient of soil melting. | 10 |
| a2 | The 2nd coefficient of soil melting. | 0.006 |
| SNr | Rate of snow melting (mm/C/day). | 3 |
| SNmt | Minimum temperature for snow melting (ºC). | 0 |
| K9 | The days over which to sum temperature to calculate start of growth. | 14 |
| K10 | The days over which to sum temperature to calculate start of soil melting. | 10 |
| b1 | The critical growth rate (Vcr or Vs). | 0.04 |
| b4 | The correction of growth rate (Gr*b4). | 0.6 |
| b5 | The correction of Vo(j*b5) and Vmin(j*b5). | 1.9 |
| b6 | Vo[j]= b6*j+b7. | 0.25 |
| b7 | b7 and b6 determine the function Vo[j]. | 0.42 |
| b8 | Vmin[j]=(EXP((b8+j)*0.4)-5.0)*b9. | 2.5 |
| b9 | b8 and b9 determine the function Vmin[j] | 0.04 |
| b10 | The growth rate in the S, G2 & M phases. | 1 |

Table S2. Site information of tree-ring width data.

| Site ID | Species | Simulation period | Lon. (º) | Lat. (º) | Elev. (m) | Detrending method |
|---|---|---|---|---|---|---|
| FIRT | *Picea glauca* | 1950–2022 | -141.63 | 68.65 | 790 | (Edwards et al., 2025) |
| QUEB | *Picea mariana* | 1950–2017 | -68.79 | 50.81 | 531 | (Wang et al., 2020, 2022) |
| TORN | *Pinus sylvestris* | 1950–2004 | 19.63 | 68.26 | 400 | (Grudd, 2008) |
| ATLA | *Cedrus atlantica* | 1950–2001 | -5.07 | 32.96 | 2200 | (Esper et al., 2007) |
| JING | *Pinus tabulaeformis* | 1950–1999 | 104.7 | 36.6 | 2350 | (Yang et al., 2023) |
| WHIT | *Pinus longaeva* | 1956–1977 | -118.17 | 37.45 | 3200 | (Salzer et al., 2009) |
| TIAN | *Pinus taiwanensis* | 1950–1999 | 119.33 | 30.33 | 1100 | (Wang et al., 2019) |

Table S3. Comparison of CMA-ES calibrated VSM parameters at FIRT and default ones used in the MATLAB version. The calibrated VSM parameters were the median of the 100 best simulations in each CMA-ES experiment.

| **Symbol** | **C5000** | **C10000** | **C15000** | **C20000** | **C30000** | **C50000** | **Default** |
|---|---|---|---|---|---|---|---|
| Tf1 | 4.02 | 3.76 | 4.99 | 4.73 | 4.03 | 3.80 | 3 |
| Tf2 | 19.53 | 19.75 | 12.02 | 15.58 | 17.20 | 18.50 | 16 |
| Tf3 | 20.22 | 20.96 | 23.45 | 23.31 | 22.23 | 24.34 | 22 |
| Tf4 | 30.75 | 36.30 | 32.52 | 28.96 | 37.65 | 30.83 | 30 |
| Wf1 | 0.02 | 0.03 | 0.02 | 0.01 | 0.01 | 0.02 | 0.01 |
| Wf2 | 0.26 | 0.14 | 0.06 | 0.31 | 0.20 | 0.19 | 0.05 |
| Wf3 | 0.81 | 0.75 | 0.79 | 0.79 | 0.81 | 0.80 | 0.8 |
| Wf4 | 0.93 | 0.91 | 0.95 | 0.96 | 0.90 | 0.88 | 0.9 |
| SNo | 233.10 | 262.38 | 449.80 | 177.17 | 237.50 | 241.46 | 300 |
| Wo | 0.11 | 0.13 | 0.00 | 0.30 | 0.39 | 0.09 | 0.2183 |
| Wmax | 0.24 | 0.32 | 0.22 | 0.42 | 0.30 | 0.27 | 0.7 |
| Wmin | 0.05 | 0.05 | 0.03 | 0.03 | 0.03 | 0.04 | 0.01 |
| rootd | 519.27 | 598.62 | 901.81 | 463.69 | 1088.17 | 706.10 | 500 |
| rated | 0.05 | 0.08 | 0.09 | 0.05 | 0.03 | 0.06 | 0.0051 |
| Pmax | 25.34 | 14.16 | 37.51 | 38.76 | 32.11 | 25.96 | 30 |
| k1 | 0.52 | 0.78 | 0.87 | 0.54 | 0.41 | 0.74 | 0.9 |
| k2 | 0.50 | 0.53 | 0.58 | 0.20 | 0.21 | 0.40 | 0.25 |
| k3 | 0.13 | 0.18 | 0.09 | 0.14 | 0.02 | 0.17 | 0.18 |
| Tm | 48.89 | 77.65 | 28.79 | 33.95 | 9.81 | 65.81 | 20 |
| Tg | 38.23 | 6.36 | 8.30 | 45.71 | 38.48 | 25.94 | 30 |
| a1 | 3.31 | 3.96 | 13.80 | 13.45 | 9.13 | 15.90 | 10 |
| a2 | 14.42 | 7.85 | 10.53 | 2.36 | 14.70 | 7.14 | 0.006 |
| SNr | 1.73 | 2.48 | 2.70 | 1.04 | 1.17 | 2.27 | 3 |
| SNmt | 2.55 | 4.70 | 4.63 | 2.37 | 2.65 | 4.34 | 0 |
| K9 | 9.82 | 10.48 | 12.74 | 8.38 | 11.66 | 13.32 | 14 |
| K10 | 9.55 | 7.86 | 8.09 | 14.98 | 12.82 | 12.85 | 10 |
| b1 | 0.37 | 0.48 | 0.44 | 0.07 | 0.20 | 0.30 | 0.04 |
| b4 | 1.84 | 2.02 | 1.37 | 1.59 | 0.83 | 1.50 | 0.6 |
| b5 | 2.61 | 1.13 | 2.71 | 0.81 | 2.00 | 2.32 | 1.9 |
| b6 | 2.17 | 1.15 | 1.50 | 0.24 | 0.52 | 1.69 | 0.25 |
| b7 | 2.84 | 2.92 | 1.11 | 4.00 | 4.89 | 1.74 | 0.42 |
| b8 | 0.06 | 2.55 | 1.19 | 3.02 | 0.40 | 0.14 | 2.5 |
| b9 | 0.66 | 0.60 | 0.79 | 0.27 | 0.30 | 0.59 | 0.04 |
| b10 | 1.90 | 5.23 | 5.27 | 2.76 | 6.67 | 1.22 | 1 |

Table S4. Comparison of CMA-ES calibrated VSM parameters at WHMT and default ones used in the MATLAB version. The calibrated VSM parameters were the median of the 100 best simulations in each CMA-ES experiment. The experiment in red showed extraordinarily high skill with Pearson's r greater than 0.9 against observed TRW.

| Symbol | C5000 | C10000 | C15000 | C20000 | C30000 | C50000 | Default |
|---|---|---|---|---|---|---|---|
| Tf1 | 9.62 | 2.56 | 0.94 | 1.04 | 7.02 | 4.92 | 3 |
| Tf2 | 19.52 | 15.23 | 17.25 | 18.85 | 17.6 | 19.22 | 16 |
| Tf3 | 25.72 | 22.12 | 22.12 | 22.70 | 23.22 | 23.46 | 22 |
| Tf4 | 38.93 | 34.86 | 31.39 | 29.34 | 31.86 | 30.86 | 30 |
| Wf1 | 0.034 | 0.029 | 0.035 | 0.037 | 0.028 | 0.028 | 0.01 |
| Wf2 | 0.12 | 0.19 | 0.23 | 0.21 | 0.078 | 0.28 | 0.05 |
| Wf3 | 0.80 | 0.74 | 0.75 | 0.80 | 0.77 | 0.76 | 0.8 |
| Wf4 | 0.94 | 0.93 | 0.95 | 0.90 | 0.93 | 0.85 | 0.9 |
| SNo | 67.5 | 44.83 | 27.95 | 338.55 | 22.75 | 42.12 | 300 |
| Wo | 0.14 | 0.30 | 0.188 | 0.26 | 0.32 | 0.37 | 0.2183 |
| Wmax | 0.43 | 0.50 | 0.25 | 0.37 | 0.37 | 0.211 | 0.7 |
| Wmin | 0.012 | 0.0065 | 0.014 | 0.033 | 0.023 | 0.014 | 0.01 |
| rootd | 215.86 | 245.16 | **659.96** | 430.46 | **1725.98** | **978.23** | 500 |
| rated | 0.0126 | 0.0758 | 0.018 | 0.06 | 0.014 | 0.017 | 0.0051 |
| Pmax | 6.35 | 48.58 | 16.42 | 32.63 | 12.20 | 33.45 | 30 |
| k1 | 0.675 | 0.64 | 0.93 | 0.96 | 0.44 | 0.66 | 0.9 |
| k2 | 0.29 | 0.94 | 0.53 | 0.43 | 0.57 | 0.34 | 0.25 |
| k3 | 0.23 | 0.24 | 0.03 | 0.77 | 0.016 | 0.11 | 0.18 |
| Tm | 20.24 | 7.67 | 54.00 | 50.67 | 21.69 | 24.60 | 20 |
| Tg | 29.43 | 15.49 | 45.15 | 48.24 | 46.28 | 25.46 | 30 |
| a1 | 14.17 | 12.57 | 6.94 | 17.95 | 8.62 | 17.50 | 10 |
| a2 | 6.29 | 5.03 | **10.74** | 6.13 | **13.23** | **15.13** | 0.006 |
| SNr | 8.27 | 7.26 | 5.64 | 6.15 | 6.21 | 8.30 | 3 |
| SNmt | 2.37 | 3.63 | 9.11 | 1.28 | 9.48 | 8.65 | 0 |
| K9 | 14.39 | 9.22 | 14.15 | 14.70 | 9.68 | 12.52 | 14 |
| K10 | 10.84 | 10.10 | 8.02 | 13.59 | 14.43 | 8.86 | 10 |
| b1 | 0.30 | 0.24 | 0.14 | 0.22 | 0.15 | 0.24 | 0.04 |
| b4 | 4.28 | 2.51 | 1.04 | 4.09 | 1.11 | 1.18 | 0.6 |
| b5 | 2.58 | 2.62 | 1.71 | 1.43 | 2.45 | 3.01 | 1.9 |
| b6 | 0.78 | 2.25 | 2.05 | 2.88 | 2.66 | 2.39 | 0.25 |
| b7 | 2.82 | 3.62 | 2.78 | 0.95 | 0.98 | 3.81 | 0.42 |
| b8 | 1.36 | 0.48 | 1.61 | 0.049 | 0.84 | 0.76 | 2.5 |
| b9 | 0.15 | 0.55 | 0.88 | 0.77 | 0.72 | 0.21 | 0.04 |
| b10 | 3.33 | 6.04 | 4.23 | 2.67 | 1.00 | 1.79 | 1 |
